# Fluorescence Characterization of Extracellular Vesicles using Single-Molecule Confocal Microscopy

**DOI:** 10.1101/2025.04.21.649767

**Authors:** Tianxiao Zhao, Noelia Pelegrina-Hidalgo, Daniel C. Edwards, Krzysztof M. Bąk, Utsa Karmakar, Anuruddika J. Fernando, Marc Vendrell, Adriano G. Rossi, Scott L. Cockroft, Tilo Kunath, Rebecca S. Saleeb, Mathew H. Horrocks

## Abstract

Extracellular vesicles (EVs) are small, membrane-bound particles released by cells into the extracellular environment. They play a pivotal role in cell communication and have recently gained prominence as biomarkers. However, their low abundance and high heterogeneity challenges their accurate characterization using conventional approaches. To enable the specific detection of individual EVs, we coupled EV-specific antibodies labeled with two different fluorophores with fast-flow microfluidics and single-molecule confocal microscopy. This allowed us to determine the concentration of EVs down to femtomolar levels (∼10^7^ EVs/mL), and we demonstrated the approach’s capacity to detect EVs even in the presence of other lipid vesicles. We highlighted the ability to quantify EVs in serum and plasma samples, without the need for purification. Furthermore, we compared the yield of EVs extracted from both serum and plasma using ultracentrifugation and various size exclusion chromatography approaches. Overall, our method offers a highly specific, sensitive and easy-to-use solution for characterizing EVs from different sources.

## Introduction

Extracellular vesicles (EVs) are small lipid vesicles that are naturally secreted by all cell types.^[1]^ They play important roles in biological processes, such as intercellular communication and clearance of unwanted biomolecules.^[2]^ EVs reflect their cellular origin through distinct surface markers and specific cargoes, including proteins, nucleic acids and other biomolecules.^[3]^ These features make them useful for studying cell-specific processes and disease states, acting as a “window” into the originating cells and their physiology.^[3]^ For this reason, EVs are being explored as biomarkers for cancers, infectious diseases, and neurological disorders.^[4–8]^

Characterizing EVs presents challenges due to their low abundance and variability in size and surface markers, and the presence of other similar sized particles in biological samples. Traditional biochemical techniques struggle with these issues, and so analysis often occurs at the single-EV level. Detecting EVs, which are about one hundred times smaller in diameter than cells, requires sensitivity four orders of magnitude higher than conventional flow cytometry. Recent studies have shown EVs may contain fewer than ten of each surface protein,^[9]^ necessitating single-molecule methods for accurate quantification. Techniques such as nanoparticle tracking analysis (NTA) can size and quantify vesicles accurately, but their reliance on light scattering limits specificity, and they cannot distinguish EVs from similarly sized particles such as lipoproteins.^[10]^ These impurities increase sample heterogeneity and risk inaccurate measurements.^[11]^

Recent methods to address EV heterogeneity and enhance specificity include nano-flow cytometry,^[12]^ and microscopic imaging of individual EVs^13,14,14–17]^. While nano-flow cytometry provides specificity, it relies on costly equipment. Microscopy, on the other hand, is time-consuming and requires specialized surfaces to enhance immobilization and reduce background signal.^[13]^ Furthermore, microscopy approaches are limited to imaging only the EVs present in each field of view, typically <10^4^, and so are low throughput, limiting the detection of rare populations.

Single-molecule confocal (smConfocal) microscopy provides a sensitive alternative to surface-based approaches, while also enabling the sampling of heterogeneous populations.^**[18,19]**^ We have recently used smConfocal microscopy with fast-flow microfluidics^**[18]**^ to both measure the ability of biomolecules to permeabilize membranes,^**[20]**^ and to detect antibody-bound protein aggregates formed during Parkinson’s disease.^**[21]**^ The latter involved using equimolar mixtures of the same antibody tagged with either of two orthogonal fluorophores, which were detected using Two-Color Coincidence Detection (TCCD).^**[22]**^ Unbound antibody and monomeric protein appeared single-colored, whereas oligomers, having many of copies of the same epitope, could bind multiple antibodies, and so were two-colored.

In the work presented here, we introduce Vesicle Imaging by Single-molecule TCCD Analysis (VISTA), an innovative approach that achieves highly specific detection of EVs at femtomolar concentrations, directly in biofluids. By using antibodies labeled with two orthogonal fluorophores, fast-flow microfluidics and single-molecule confocal microscopy, VISTA surpasses current methods in specificity, distinguishing EVs from other similar-sized particles in complex samples. Besides higher specificity, VISTA compares favorably with current characterization techniques with regards to sensitivity. This easy-to-use approach streamlines EV analysis without requiring purification, offering significant potential for biomarker discovery and disease diagnostics, making VISTA an essential tool for EV characterization.

## Results and Discussion

### Sensitive single-molecule detection of extracellular vesicles

To specifically detect individual EVs, we labeled two populations of the same antibody targeting CD9, a tetraspanin surface EV marker commonly used for EV characterization,^[3]^ with the orthogonal fluorophores Alexa Fluor 488 (AF488) and Alexa Fluor 647 (AF647). As free protein will only contain a single epitope, they will be bound by either an AF488- or AF647-labeled antibody. EVs, on the other hand, have more than one CD9 molecule on their surface^[23]^, and therefore bind both AF488- and AF647- labeled antibodies (***Figure 1A***). EVs therefore give rise to coincident fluorescent bursts from both fluorophores as they transit the diffraction-limited confocal volume (***Figure 1B***).

**Figure 1.**
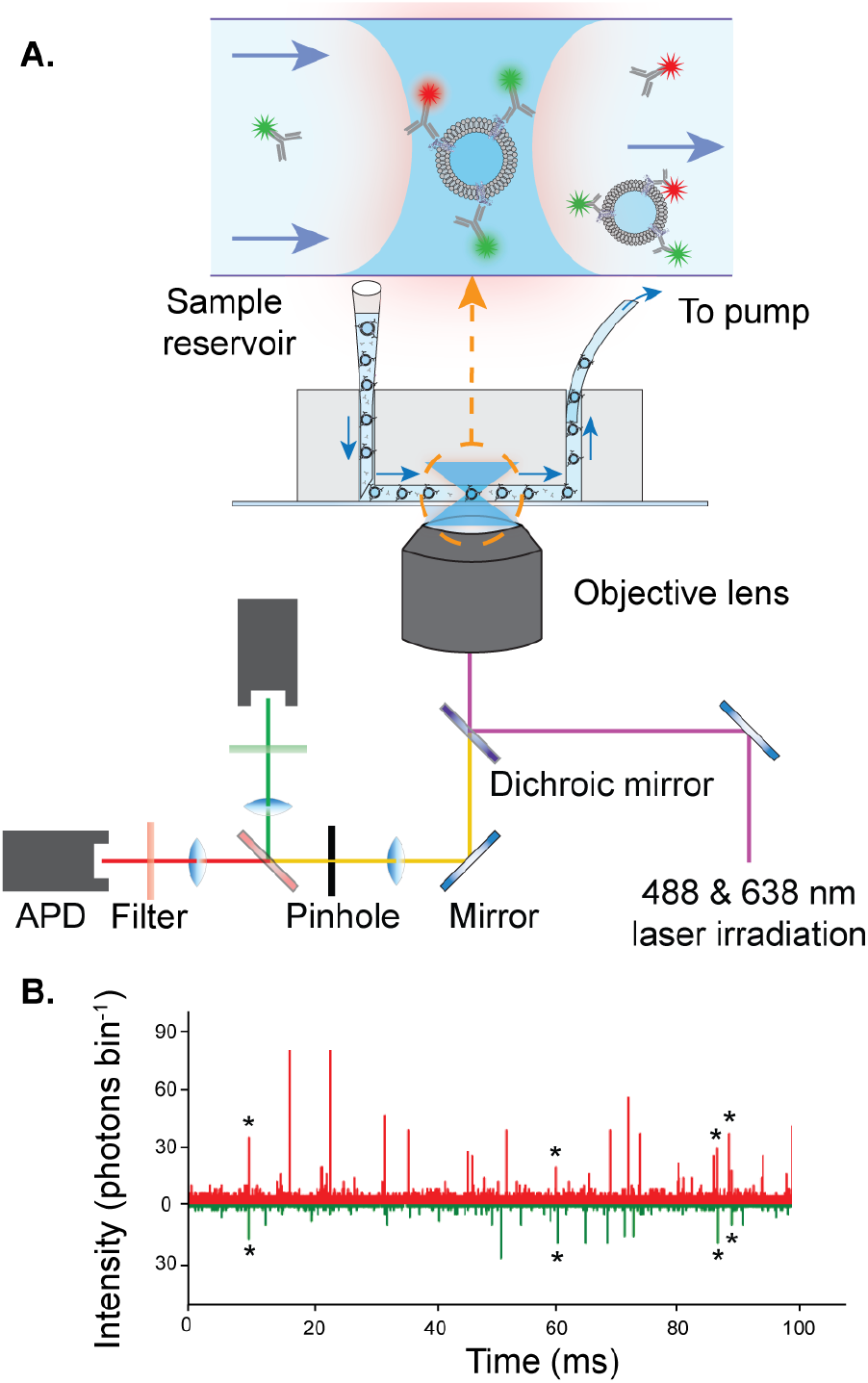
Vesicle Imaging by Single-molecule TCCD Analysis (VISTA) A) Schematic of the experimental setup for EV detection using a fast-flow microfluidics and single-molecule confocal microscopy. B) Representative TCCD single-molecule data showing EV detection with AF488-tagged antibodies (green) and AF647-tagged antibodies (red). Stars show example coincident bursts corresponding to dual antibody-tagged EVs.

We first sought to assess the sensitivity of our approach for detecting EVs. This was achieved by preparing solutions of EVs originating from a mammalian cell line (HCT116), with concentrations spanning several orders of magnitude into a solution containing AF488- and AF647-tagged antibody. Both the event rate (***Figure 2A***), defined as the number of CD9-positive EVs per unit time, and the association quotient (Q), which is a measure of the fraction of coincident events (for further details, see ***Supporting Information***) increased at higher EV concentrations (r^2^= 0.9984 and r^2^=0.9741, respectively) (***Figure 2B***). To convert the association quotient to an EV concentration, we made use of dye-filled synthetic vesicles as a calibrant (see ***Supporting Information*** and ***Figure S1***)

**Figure 2.**
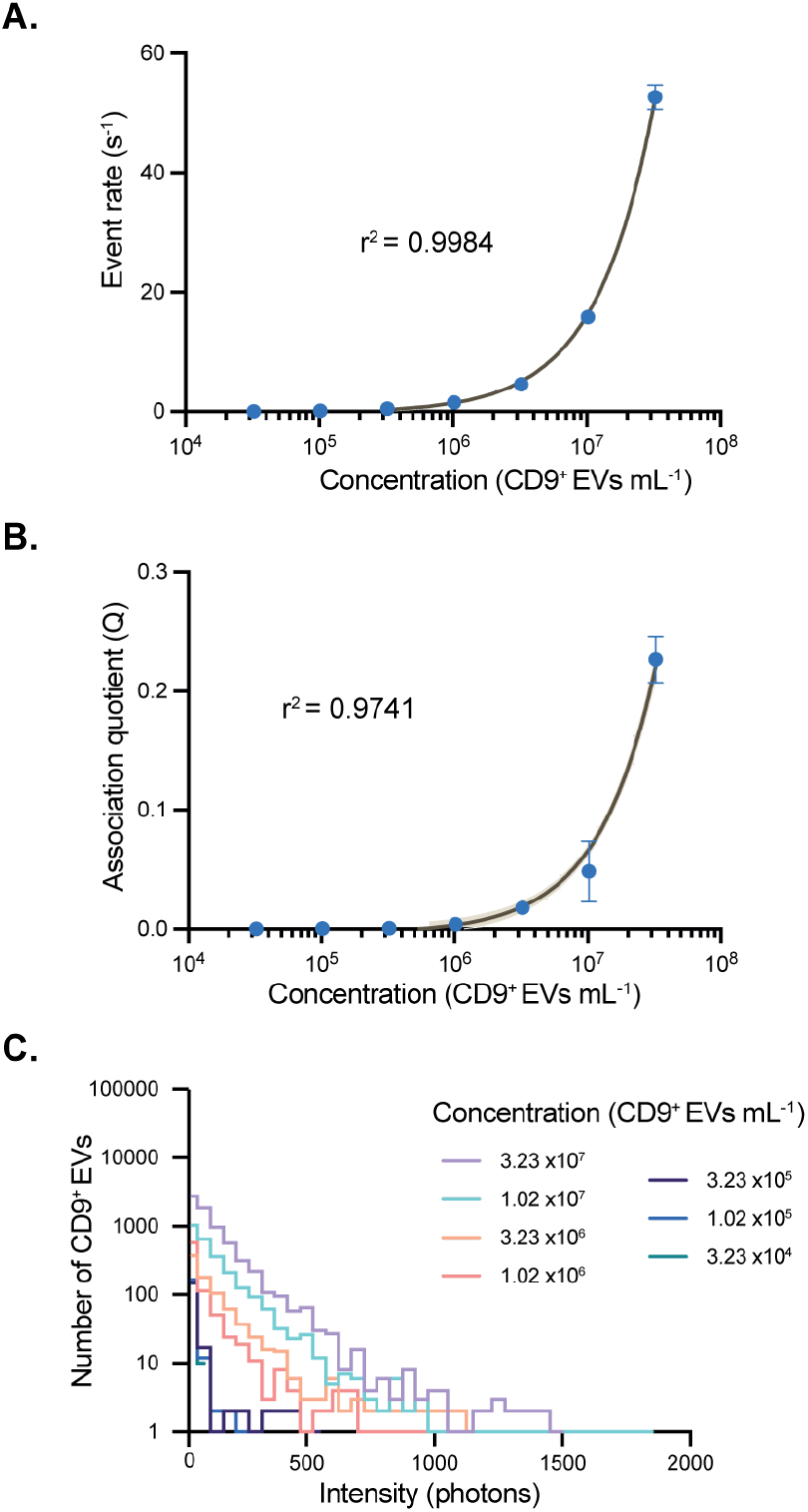
VISTA measurement of a concentration series of EVs with fixed concentration of antibodies. A) Event rate of coincident bursts and B) association quotient (Q) shown for increasing concentrations of EVs. C) Intensity histograms for each EV concentration measured in A) and B). Data are shown as mean ± SD, n = 3, the shaded band is a pointwise 95% confidence interval on the fitted values.

In addition to quantifying the number of EVs, the intensity of individual tagged EVs was determined, which is proportional to the number of bound antibodies. Example intensity histograms for a range of EV concentrations are presented in ***Figure 2C***. At an EV concentration of 3.23 x 10^7^ particles /mL, the mean intensity of the tagged EVs was 132.13 photon counts bin^-1^, which corresponds to ∼3 antibodies per EV (see ***Supporting Information*** for details of estimation). Stoichiometry histograms for a range of EV concentrations are shown in ***Figure* S2**, and mean intensities for all concentrations of EVs are presented in ***Table S1***.

To quantify the sensitivity of our approach, we determined both the limit of blank (LoB), and the limit of detection (LoD) from a sample containing no analyte, and one with a low concentration of EVs. Utilizing the association quotient as our readout, we calculated a LoB of 5.6 x10^5^ CD9^+^ EVs mL^-1^ (0.93 fM) and a LoD of 5.7 x10^5^ CD9^+^ EVs mL^-1^ (0.95 fM) for CD9^+^ EV detection using VISTA. This compares favorably with other common EV characterization approaches, such as NTA using ZetaView, which has an LOD of 1 x10^5^ particles mL^-1^. We also analyzed the same EV samples using a surface-based single-EV detection approach commercially available from Oxford Nanoimaging, determining an LOD of 4 x 10^8^ EVs mL^-1^ (∼600 fM) (***Figure S3*** and ***Supporting Information***).

As it is not always possible to obtain or directly label antibodies, we also demonstrated that VISTA could be performed using fluorophore-tagged secondary nanobodies targeting unlabeled antibodies, according to our recently optimized protocols^[24]^ (***Figure S4*** and ***Supporting Information*)**.

The high sensitivity of VISTA is a result of two factors. Firstly, maintaining a low concentration of the antibody allows for the observation of individual molecules passing through the confocal volume. Secondly, fast-flow microfluidics enables a rapid data acquisition rate, increasing the throughput of detected and characterized events over a short acquisition time.

### Specificity of VISTA for extracellular vesicles

Approaches commonly used for quantifying EVs, such as NTA, rely on the scattering of light for particle detection. While these allow the concentration and size of particles to be measured, they are typically unable to distinguish EVs from other similar-sized particles, such as lipoproteins,^[10]^ which are common contaminants in purified EVs. We therefore sought to determine whether VISTA could distinguish EVs from similar-sez particles, such as large unilamellar vesicles (LUVs).

To achieve this, samples were prepared with varying EV:LUV ratios. While the concentrations of EVs were diluted to concentrations spanning several orders of magnitude, the total number of particles (EVs + LUVs) was kept constant by adjusting the concentration of LUVs (***Figure 3A***). The samples were measured using NTA and VISTA to determine particle or CD9^+^ EV concentrations, respectively. As expected, particles were detected in all samples using the ZetaView, showing an EV-independent trend, since the measured concentration did not vary as EV:LUV ratio increased (***Figure 3B***; r^2^ = 0.1432). VISTA, on the other hand, detected vesicles in an EV-dependent manner, increasing as the mixture was enriched with EVs (***Figure 3C***; r^2^ = 0.9883). Taken together, these results demonstrate that not only is VISTA as sensitive as NTA for particle detection, but it can detect EVs specifically.

**Figure 3.**
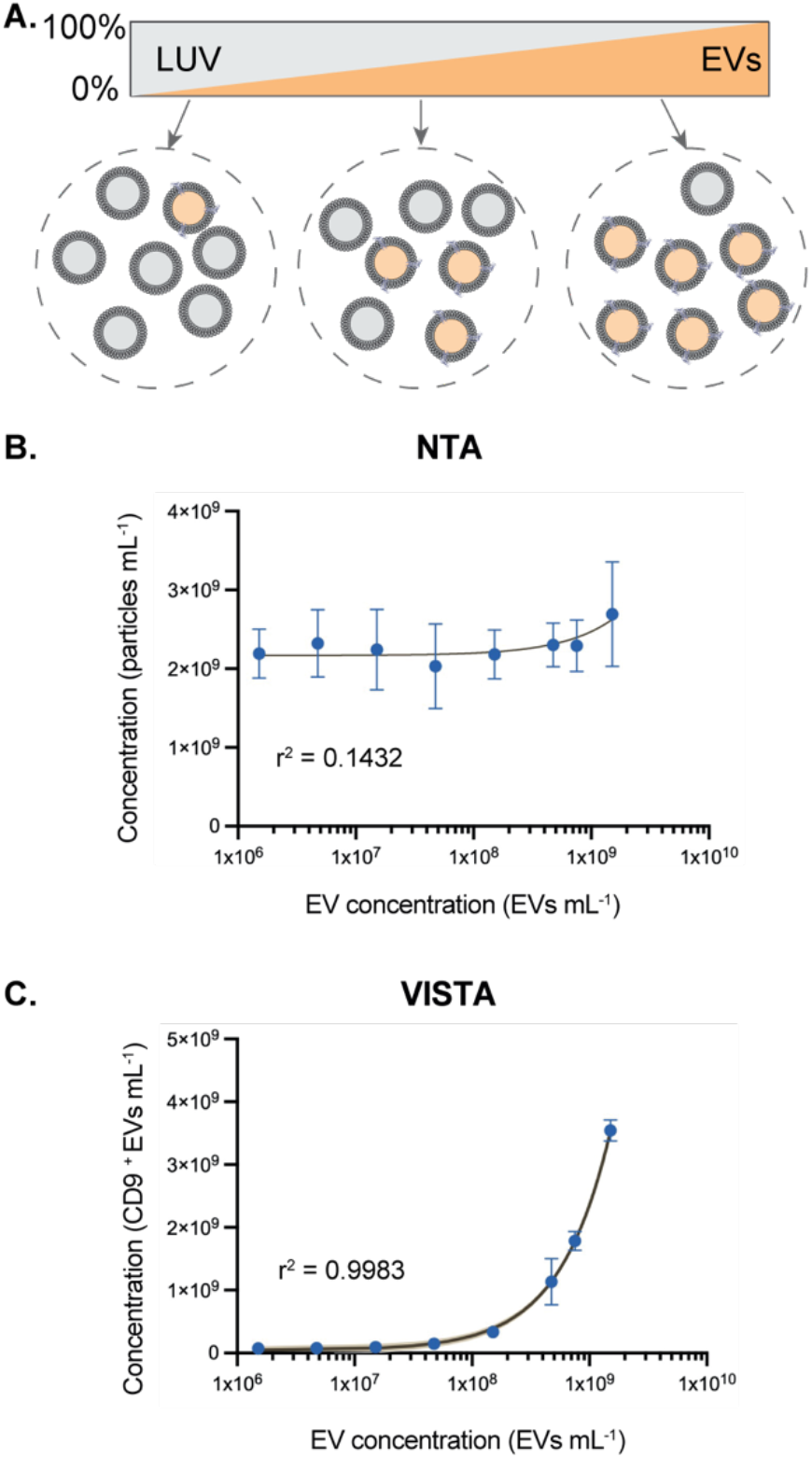
Specificity of VISTA for quantifying EVs in heterogeneous solutions. A) The number of total particles was kept constant among different samples by increasing the EV (orange):LUV(grey) ratio. B) NTA measurements of mixtures. C) VISTA measurements of samples. Data are shown as mean ± SD, n = 3, the shaded band is a pointwise 95% confidence interval on the fitted values.

### Quantifying EVs in human biofluids using VISTA

EVs have gained prominence as a source of biomarkers for various diseases, including cancer ^[25]^ and Parkinson’s disease^[26]^. Blood-derived EVs are of special interest due to the accessibility of this biofluid in the clinic. Conventional EV analysis approaches, such as NTA, require biofluids to be processed due to their heterogeneous nature. For this, EV-sized particles are isolated from biofluids using commercially available kits, size exclusion chromatography (SEC), or ultracentrifugation (UC). These extra steps can lead to disruption and/or loss of EVs, which is particularly problematic for biomarkers present at low concentration and/or limited patient sample volumes. It can also lead to limitations for accurate EV characterization. VISTA, however, can be performed directly on small volumes of biofluids (as low as 3 µL), avoiding these issues and making it a straightforward, isolation-free approach for EV quantification.

To demonstrate this, we first separated blood from three donors into either plasma or serum, and followed the same VISTA protocol as before, using AF488- and AF647-labeled CD9 antibodies for detection (***Figure 4A***). Using VISTA, we were able to detect green and red coincident bursts, showing the capacity of VISTA to measure CD9^+^ EVs from unprocessed biofluids.

**Figure 4.**
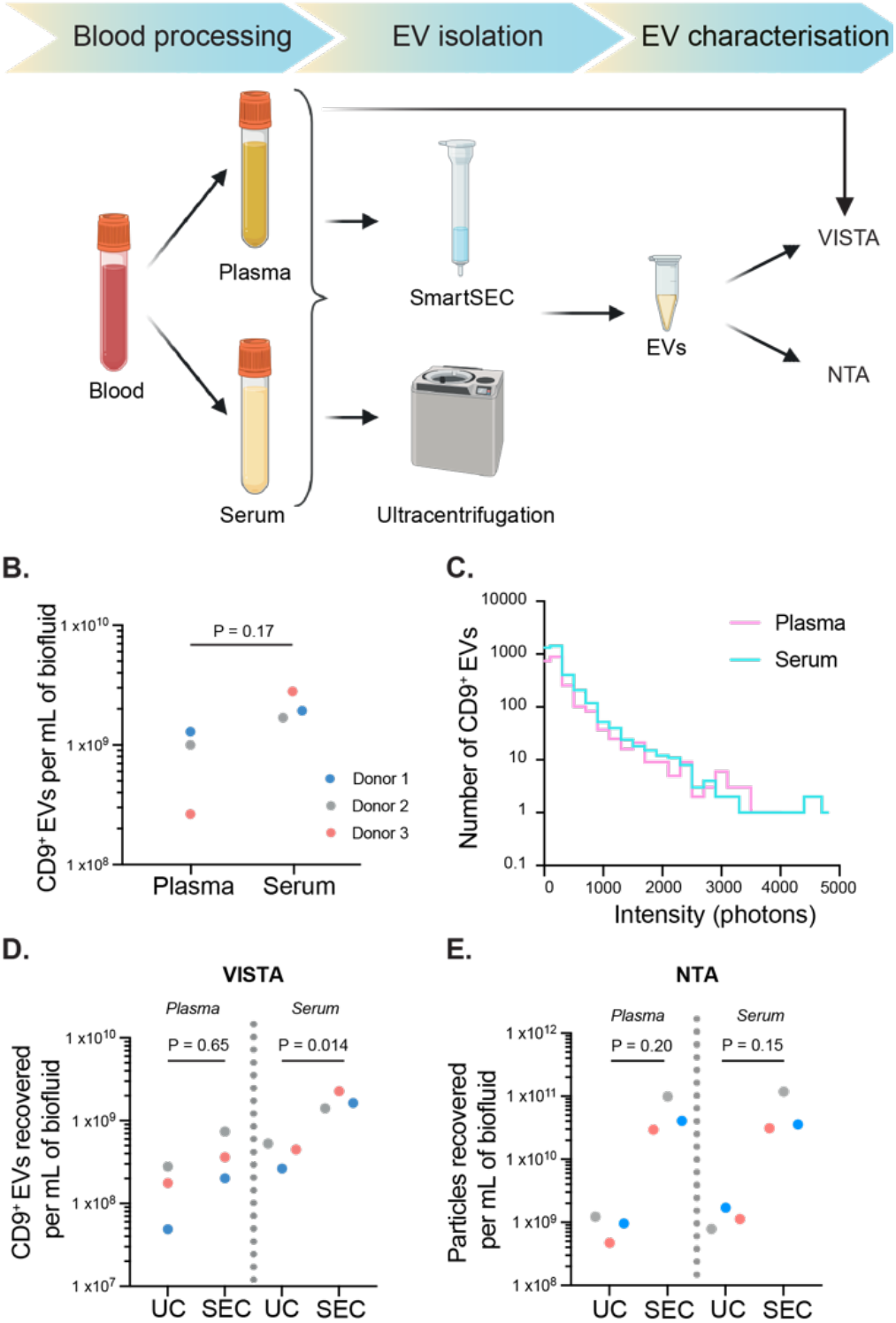
Quantifying EVs in human biofluids using VISTA. (A) Schematic showing blood processing and EV isolation methods used prior to VISTA or NTA (ZetaView) measurements. B) VISTA measurements of plasma and serum in three different donors (refer to color code for each donor). C) histogram intensity distribution is shown for both neat biofluids (blue: plasma; purple: serum). D) VISTA measurements of ultracentrifugation (UC) or SmartSEC (SEC) isolated EVs from plasma and serum. E) NTA measurements of same samples. For B), D), and E), points show results from individual donors. p-values were calculated by the Student t-test (paired).

Although VISTA did not show statistically significant differences in the levels of EVs in serum and plasma (p = 0.17; Student t-test (paired)) due to large inter-individual variance (***Figure 4B***), a trend of higher CD9^+^ EVs in serum was observed when compared within donors. This aligns with previous findings from other studies,^[27]^ likely due to an increase in platelet-derived EVs caused by the activation of platelet clotting pathways.^[28]^ We also looked at the intensity distributions of the CD9^+^ EVs (***Figure 4C, Table S2***), which showed a similar intensity distribution to CD9^+^ EVs from mammalian cells (***Figure 2C*** and ***Table S1***), demonstrating that the serum and plasma EVs contained a similar number of CD9 molecules to those from the cell lines. This shows VISTA’s advantage over NTA EV characterization, which requires EV isolation.

To study EV-enriched samples, isolation can be performed in biofluids using several alternative approaches, including UC^[29]^, and SEC^[30]^. We therefore decided to use VISTA to evaluate how different methods affect EV recovery from plasma and serum, and compared VISTA to NTA-measured concentrations. For SEC, there are numerous commercially available kits, and so we first compared these using VISTA and NTA (***Figure S5***), and found that SmartSEC gave the highest yield of EVs. We then evaluated the EV yield from SmartSEC (now referred to as SEC) and UC on both plasma and serum from three donors (***Figure 4D***). Similar to neat biofluids, the number of CD9+ EVs isolated per mL of serum was overall higher than that from plasma, regardless of the isolation protocol. Both biofluid-isolated samples consistently showed that SEC gave a higher yield of CD9+ EVs compared with UC (plasma-UC vs plasma-SEC: p = 0.65; serum-UC vs serum-SEC: p = 0.014; Student t-test (paired)).

Both SEC and UC separate particles based on their size alone, and so are not specific to EVs, leading to the co-isolation of multiple vesicle types and other similarly sized contaminants present in blood. We therefore measured the same purified samples using NTA (***Figure 4E***), showing ∼10-100-fold increase in the number of particles recovered per mL of biofluid when compared to VISTA (***Figure 4D***), due to the detection of non-CD9+ vesicles of a similar size. As with VISTA, SEC isolation led to a higher recovery of particles than UC in both biofluids (plasma-UC vs plasma-SEC: p = 0.2; serum-UC vs serum-SEC: p = 0.15; Student t-test (paired)).

## Conclusions

VISTA offers significant advantages for the detection and characterization of EVs. VISTA can specifically detect EV concentrations down to 5.7 x10^5^ EVs mL^-1^ (femtomolar range) using CD9 as an EV-specific marker. Its specificity allows for the distinction between EVs and similarly sized particles, such as lipoproteins, enhancing accuracy in complex biofluids. Additionally, VISTA allows for the identification of EV subpopulations within heterogeneous samples, providing the capacity to obtain a complete profile of EVs based on their surface markers. Importantly, VISTA facilitates the direct analysis of neat biofluids without requiring EV isolation, thus preventing potential sample loss or disruption, and streamlining the process. This method’s ability to be used with any EV surface markers allows for custom EV detection, making VISTA a powerful tool for biomarker discovery and disease research.

## Acknowledgements

We wish to thank the blood donors who contributed samples for this work, and Sonja Vermeren and her PhD students, Alfie Sanderson and Caroline Pumpe, who assisted with blood collection and processing. We also would like to thank Laila Kjellström who donated funds used towards the purchase of instruments used in this study. The single-molecule instruments used in this study were funded by the UK Dementia Research Institute, UCB Biopharmaceuticals, Alzheimer’s Research UK (ARUK-EG2018B-004) and a kind donation from Dr. Jim Love. T.Z. was supported by a Centre for Doctoral Training in Tissue Repair, Innovation and Collaboration (CenTRIC) studentship funded by the Eureka Foundation. N.P.H. was supported by a Medical Research Scotland studentship and Oxford Nanoimaging Ltd. K.M.B and D.C.E, were supported by the Leverhulme Trust Research Grant (Grant no. RPG-2021-328).

