## Supplementary Information for "Fluorescence Characterization of Extracellular Vesicles using Single-Molecule Confocal Microscopy"

### S1. Materials and Methods

#### S1.1. Antibody labeling

Anti-CD9 antibodies (100-0138, STEMCELL Technologies) were purchased and conjugated in-house with Alexa Fluor dyes using antibody labeling kits (A20181 and A20186, Invitrogen), following the manufacturer's instructions. Antibody concentrations were determined using a NanoDrop spectrophotometer.

#### S1.2. Preparation of microfluidic device

The microfluidic device design has been described previously,[1] and consists of a single channel (width = 100 µm, height = 25 µm, length = 1 cm). Microfluidic devices were fabricated using standard soft-lithography techniques into polydimethylsiloxane (PDMS; Dow Corning) with SU-8 photoresist on silicon masters, as described previously.[1] The channels were bound to glass coverslips (VWR, thickness = 1) after exposure to an oxygen plasma (ZEPTO B, Diener).

##### S1.3. Single-molecule confocal measurements

All confocal microscopy measurements were performed on a custom built single-molecule confocal microscope described previously.[1] An Oxxius L6Cc laser combiner (Oxxius, France) containing Gaussian laser beams of two wave lengths 488 nm (LBX-405-100-CSB-OE, Oxxius) and 638 nm (LBX-405-100-CSB-OE, Oxxius) was used as a laser source. Laser beams were overlapped within a 0.13 NA optical fiber (Thorlabs) and collimated by a reflective collimator (RC08APC-P01, Thorlabs), and directed through the back-port of an inverted microscope (Nikon TE2000-U). A dichroic mirror (DI03-R405/488/561/635, Semrock) reflected the beam through an oil-immersion objective lens (Nikon CFI Plan Apochromat VC 100x Oil, NA 1.4, W.D 0.13 mm) that focused the light to a confocal spot. The emitted fluorescence was collected by the same objective lens and was passed through the same dichroic mirror, before being focused by a tube lens within the microscope body through a 50 µm pinhole (Thorlabs). A second dichroic mirror (585DRLP, Omega Filters) was used to separate the fluorescence from the two different fluorophores. The longer wavelength passed through the dichroic mirror and was focused by a lens (Plano apoconvex, focal length = 50 mm, Thorlabs) through a filter set (long-pass: LP02-647RU-25, Semrock, and band-pass: FF01-432/515/595/730-25, Semrock) onto an Avalanche Photodiode (APD) detector (PerkinElmer). The shorter wavelengths were reflected and focused through a second filter set (long-pass: BLP01-488R-25 and band-pass: FF01-525/30-25, Semrock) onto a second APD. Outputs from the APDs were connected to a USB data acquisition card (USB-CTR04, Measurement Computing), which counted the signals and combined them into time-bins of 100 µs, the expected residence time of the EVs in the confocal volume. The intensities of the 488 nm and 647 nm irradiation used was 0.604 mW and 0.395 mW at the back-port, respectively.

The prepared sample was introduced into the microfluidic device using a suitable gel-loading tip. Fluidic propulsion through the channel was achieved by applying negative pressure at the distal end, facilitated by a syringe (Injekt®-F Luer Solo, B.Braun). This syringe was operated by a precision syringe pump (LEGATO 101, World precision instrument), which maintained a controlled flow rate of 96 µL $h^{-1}$, corresponding to a flow velocity of 1 cm $s^{-1}$ in the channel.

##### S1.4. Data analysis

A custom-written Python code was used to analyze all VISTA data (https://doi.org/10.5281/zenodo.15173917). The burst intensities in the green ($I_G$) and red ($I_R$) channels were first corrected for autofluorescence, calculated using the mean signal intensity in absence of fluorescent molecules (***Equations 1 and 2***). 3% of signal intensity in the red channel was determined to be cross talk from the green channel, whilst crosstalk in the other direction was negligible.

$$I_G = G - A_G \qquad \textbf{\textit{Equation 1}}$$

Where $G$ is the original intensity and $A_G$ is the autofluorescece in the blue channel (0.6 photons bin$^{-1}$).

$$I_R = R - A_R - (C \times G) \qquad \textbf{Equation 2}$$

Where $I_R$ is the modified intensity in the red channel, $R$ is the original intensity, $A_R$ is the autofluorescence in the red channel (1.97 photons bin$^{-1}$) and $C$ is the cross-talk from the green to red channel.

Simultaneous events in both channels were assessed using the AND criterion, which accepts only those signals for which both the blue-excited channel and the red-excited channel are above their respective threshold value. Thresholds were determined according to previously established threshold selection approaches.[2] To account for coincident events that could occur due to chance, a desynchronisation approach was used.[3] Time-bins in the red channel were randomly shuffled and the number of simultaneous events in the two channels above the threshold was recalculated, the resulting number of events were classified as chance events. The fraction of coincident events, with respect to chance was then calculated.

The association quotient (Q), describes the fraction of coincident events according to ***Equation 3***.

$$Q = \frac{\gamma - \delta}{\alpha + \beta - (\gamma - \delta)} \qquad \textbf{Equation 3}$$

Where $\alpha$ is the number of bursts in the green channel, $\beta$ is the number of bursts in red channel, $\gamma$ is the simultaneous events in both channels, and $\delta$ is the simultaneous events from desynchronised data.

##### S1.5. Preparation of cell-line derived EVs

Lyophilized microvesicles derived from the human colonic carcinoma cell line, HCT116 (HVM-mvHCT50, Hansabiomed), were reconstituted following the manufacturer's instructions, aliquoted into 20 µL aliquots, and stored at –80 °C. To minimize degradation, samples were subjected to no more than one freeze–thaw cycle. A stock solution containing a mixture of AF488- and AF647-conjugated anti-CD9 antibodies was prepared at a concentration of 2.5 nM. An aliquot of 2 µL of this antibody stock solution was incubated with 3 µL of freshly thawed EVs at room temperature for 1 hour. The incubation mixture was then diluted 80-fold prior to analysis. The concentration of EVs in the sample was determined using VISTA and the concentration calibration curve.

##### S1.6. Preperation of LUVs

Vesicles were prepared as described below using an Avestin LiposoFast Liposome Factory, equipped with polycarbonate membranes with 200 nm pores. Vesicle purification was carried out using PD-10 desalting columns (GE Healthcare) prepacked with Sephadex G 25 medium. All buffers and stocks solutions prepared using biological grade water (Fisher Bioreagents) and filtered with 0.22 µm syringe filters (Merck). POPC lipid (PC 16:0/18:1) and cholesterol were obtained from Sigma Aldrich.

A thin film of lipid was prepared by evaporating a deacidifed chloroform solution of POPC and cholesterol (8:2) under reduced pressure on a rotavap with high rotation to provide aliquots of 6 µmol. Films were further dried for 6 hours under high vacuum and stored at –20°C until use. Lipid films were warmed to room temperature and mixed with PBS buffer. For dye-filled calibrant LUVs, AF488- and AF647-taged strevtavidin (2.5 µM) was added to the buffer. The films were then hydrated by sonication for 30 s and vortexing for 1 hour. The lipid suspensions were then subjected to 10 freeze-thaw cycles using liquid nitrogen and a water bath (40 °C). Vesicles were then diluted and extruded 29 times through a polycarbonate membrane (pore size 200 nm). Extra-vesicular components were removed by size exclusion chromatography on a PD-10 sephadex G-25 column washed with prepared buffer (PBS). For Streptavidin conjugate-dye containing liposomes residual unincorporated protein was removed using an Izon-SEC column.

To maintain a constant total particle concentration in the detection solution, the initial concentrations of EVs and LUVs were measured by VISTA or NTA, respectively. An EV stock solution was then prepared at $2.01 \times 10^{11}$ particles $mL^{-1}$ (333.3 pM) and subjected to a half-log dilution spanning three orders of magnitude, as well as a 1:2 dilution in PBS. The total particle concentration, defined as the combined concentrations of EV and LUV, was subsequently adjusted by adding the appropriate amount of LUV.

##### S1.7. Determine the LoD and LoB of VISTA

The limit of blank (LoB), which indicates the highest number of counts expected when no analyte is detected, was calculated as:

$$LoB = mean_{blank} + 1.645(SD_{Blank})$$

where the blank sample contained no EV. The limit of detection (LoD), which denotes the lowest analyte concentration likely to be reliably distinguished from the LoB, was subsequently calculated as:

$$LoD = LoB + 1.645(SD_{lowest\ concentration})$$

The lowest concentration sample corresponds to $3.2 \times 10^4$ $CD9^+$ EVs $mL^{-1}$.

##### S1.8. Characterization of EVs using Nanoparticle Tracking Analysis (NTA)

EVs were prepared for NTA by diluting them 1:5000 or 1:1000 in 0.22 µm-filtered (Whatman) PBS (20012027, ThermoFisher Scientific). All NTA readings were measured using the Zetaview PMX-430-Z-QUATT instrument (Particle Metrix, Inning am Ammersee, Germany) and the Zetaview (version 8.05.16 SP7) software. Calibration was done using beads (provided by Particle Matrix) diluted 1:250000 (v:v) in distilled water. Measurements were performed using the 488 laser in scatter-mode with the following parameters: camera sensitivity: 65, camera shutter: 100 ms, camera frame-rate: 30, minimum trace length: 15. Cell temperature was maintained at 25°C. A total of 11 positions were recorded for each measurement. Analysis was performed using the same software, applying the following parameters: minimum brightness: 30, maximum brightness: 1000, minimum area: 10.

##### S1.9. Isolation of EVs from blood

Blood collection and processing into plasma and serum

Whole blood was collected from healthy volunteers following informed consent and ethical approval (21-EMREC-041) at the University of Edinburgh. Blood collection and processing was performed as it follows: 40 mL of freshly drawn blood was collected into tubes containing 4 ml of 3.8% sodium citrate (25116, Sigma) and centrifuged at 350 xg for 20 min at room temperature. The supernatant (~15 ml plasma) was collected and split into two vials, one for plasma-EV isolation and another for serum-EV isolation. For serum-EV isolation, 10 mL of plasma were incubated for 1.5-2h at 37 °C (water bath) with 220 µL of 1M calcium chloride to achieve platelet coagulation. Post-incubation, supernatant (serum) was transferred to a new vial, avoiding the platelet plug formed during coagulation. Then, both plasma and serum were centrifuged at 2,000 xg for 20 min to remove any debris and the supernatant was used for EV isolation (see next section). For detection of EVs in biofluids (Figure 4B & C),1 mL aliquots of each biofluid were stored at –80°C prior to VISTA.

EV isolation from plasma and serum

All isolations were performed from 1 mL of biofluid sample (plasma or serum). For all concentration steps, 100 kDa centrifugal filters (UFC503024, Merck) were used, previously rinsed with 0.22 µm-filtered Dulbecco's phosphate-buffered saline (fPBS) (14190144, ThermoFisher). Post-isolation, fresh samples were kept on ice until NTA measurements for fresh EV concentration calculations or frozen into 10 µL aliquots for frozen-thawed EV concentration calculations and storage. See below for separate methods on each isolation protocol.

*Izon column EV isolation*

Samples were concentrated to 500 µL using pre-rinsed 100 kDa filters following rounds of centrifuging at 10,000 xg for 10 min. For EV isolation, qEVoriginal columns (ICO-70, Izon Science LTD) were used, automated with the Automated Fraction Collector (AFC-V2, Izon Science LTD). Collection was performed into 400 µL fractions, and the first 4 fractions were pooled together using pre-rinsed 100 kDa filters to a final volume of 100 µL solution.

*ExoQuick Total Exosome Isolation*

Samples were processed using Total Exosome Isolation Kit (from plasma;4484450, ThermoFisher) or Total Exosome Isolation Kit (from serum; 4478360, ThermoFisher) to isolate EVs from plasma or serum, respectively. Isolations were performed following manufacturer's protocol for each biofluid type. Note plasma and serum isolations differ in the incubation times and steps to follow. Pellets were resuspended in 200 µL fPBS and mixed well, first by pipetting and then vortexing.

*SmartSEC isolation*

Samples were concentrated to 250 µL using pre-rinsed 100 kDa filters following rounds of centrifuging at 10,000 xg for 10 min. SmartSEC (SSEC200A-1, Systems Biosciences) columns were used, following the manufacturer's protocol and incubating column with sample for 30 min.

#### S2. Concentration calibration

To convert the association quotient (Q) from VISTA measurements into an absolute particle concentration, it was necessary to calibrate the measurements with samples having known concentrations. To achieve this, a solution of LUVs loaded with equal concentrations of AF488 and AF647 (5 dye molecules per LUV) to mimic antibody-bound EVs was prepared, and its concentration measured using NTA. We then performed VISTA across a wide concentration range, determining Q for each sample (***Figure S1****)*. The data were fit to a function of the non-linear regression, which was used to convert Q to concentration for VISTA EV measurements.

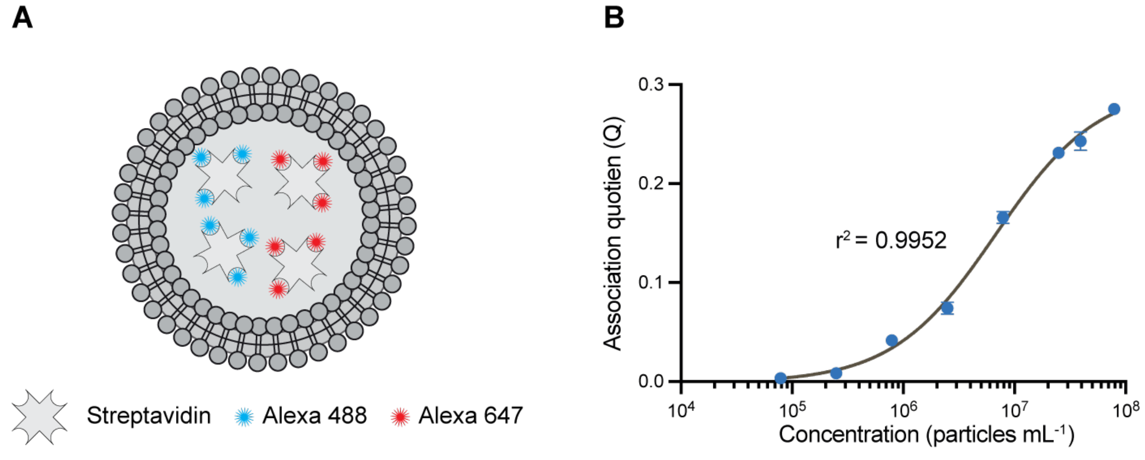

***Figure S1. VISTA measurement of dye-filled LUVs .*** *A) chemically synthesized LUVs containing AF488- and AF647-tagged streptavidin****.*** *B) The association quotient (Q) shown for varying concentrations of dye-filled LUVs. Data are shown as mean ± SD, n = 3, the shaded band is a pointwise 95% confidence interval on the fitted values. The line shows the fit of the logarithmic model:* $y = 0.0435\ln(x) - 0.5256$.

#### S3. Intensity distributions of EVs

The total intensity of each EV detected was calculated by summing the intensities from both channels. The mean intensity for each experimental condition is reported in ***Table***

***S1***. From these, the number of antibodies bound to each EV can be approximated by dividing the total intensity by the mean intensity of AF488- and AF647-tagged antibodies alone. ***Figure S2*** shows the antibody stoichiometry for EVs from the HCT116 cell line.

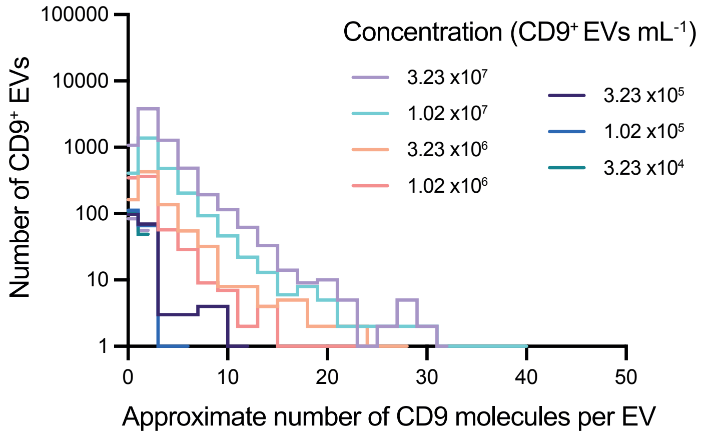

***Figure S2. Antibody stoichiometry for varying concentrations of EVs measured using VISTA.***

***Table S1 – mean intensities of EVs from HCT116 cell line.***

| **EV concentration (particles mL$^{-1}$)** | **1.51x $10^9$** | **7.53x $10^8$** | **4.76x $10^8$** | **1.5x $10^8$** | **4.76x $10^7$** | **1.51x $10^7$** | **4.76x $10^6$** | **1.51x $10^6$** |
|---|---|---|---|---|---|---|---|---|
| **Average intensity (photons)** | 132.13 | 155.18 | 136.14 | 129.40 | 85.00 | 64.48 | 48.18 | 45.02 |
| **Antibody stochiometry (number of ABs)** | 2.87 | 3.37 | 2.96 | 2.81 | 1.85 | 1.40 | 1.05 | 0.98 |

### S4. Detection of EVs using TIRF microscopy

HCT116 EVs were immobilized on microfluidic glass slides using the EV Profiler 2 (ONI) protocol and reagents. Briefly, the provided chip was incubated with Surface Reagent for 10 min, then washed and further incubated with Capture Reagent (Tetraspanin Trio) for 15 min. After washing, EVs at three different concentrations in PBS (1x$10^9$, 3.16x$10^9$ and 1x$10^{10}$ EVs/mL) and a blank (PBS only) were added to the chip and incubated for 75 min. Then, lanes were washed and fixed using Fixative reagent for 10 min. Again, lanes were washed and Staining Buffer was added and incubated for 10 min, followed by Detection reagent for 50 min. Another wash was performed and all lanes were incubated with Fixative for 5 min. Finally, all lanes were washed and the inlet/outlet holes were sealed to avoid evaporation. A total of three chips were used for 3 technical replicates.

All incubations were performed on a rocker set to 20 r.p.m., except the fixation steps, and all washes were performed using Wash Buffer provided in the kit.

Imaging was perfomed using an ONI Nanoimager equipped with a 100×/1.4 numerical aperture oil immersion objective lens and ORCA-Flash 4.0 V3 scientific complementary metal-oxide semiconductor camera. Samples were exposed sequentially to 638-, 561-, and 488-nm excitation by total internal reflection using a 53.5° illumination angle for visualization of CD81, CD63, and CD9, respectively. The resultant emission was split with a 640-nm dichroic, projecting emission above and below this wavelength onto different regions of the camera chip for dual-channel viewing. Each field of view (FOV) was imaged for 20 frames at an exposure of 30, and a 10 × 10 grid of 100-µm–spaced FOVs was captured per condition.

To quantify triple-coincident EVs ($CD9^+$ & $CD63^+$ & $CD81^+$), a coincidence analysis Fiji plugin, ComDet v.0.5.5, was used (https://github.com/ekatrukha/ComDet) by running the “Detect particles” option. Three-color events with a center of mass within 3 pixels were considered-coincident, with SD set to 4 for the 647 and 488 channels, and 4 for the 561 channel. The number of triple-coincident events was obtained for each FOV from the summary output table column “Colocalized_ch1&ch2&ch3” after merging all three channels and converting the merge into 16-bit. Data was plotted by first averaging all FOVs (FOV=100) per replicate (rep=3), and then mean of the three replicates was calculated with standard deviation. The Fiji macro used for this analysis can be found in 10.5281/zenodo.15254469

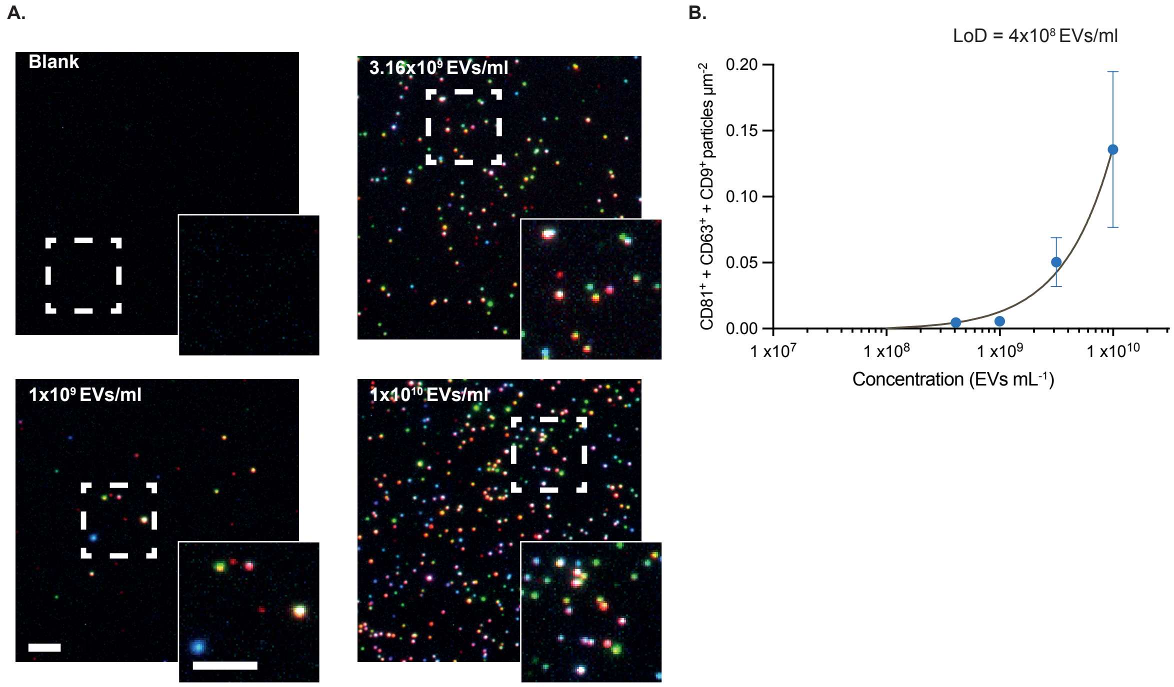

***Figure S3. HCT116-derived EVs imaging using EV Profile Kit (ONI).*** *(A) Representative images of each EV concentration used in this surface imaging assay (scale bars: 500nm). (B) Quantification of triple coincident EVs ($CD81^+$+ $CD63^+$+ $CD9^+$) per field of view. Data points show mean ± SD (n = 3 replicates, 100 FOVs/rep).*

#### S5. Detection of EVs using nanobody-tagged antibodies

It is sometimes not possible to obtain labeled antibodies for fluorescence measurements. Here, we showcase two key advantages of VISTA in detecting EVs. First, VISTA enables determination of EV origin by identifying cell-specific surface markers. EVs isolated from the LnCAP cell line were identified through their expression of the PSMA marker. Second, we demonstrate the platform's robust compatibility with nanobody-tagged antibodies by employing nanobody-conjugated PSMA and CD33 antibodies. This approach not only differentiates LnCAP-derived EVs from those expressing minimal amounts of CD33, but also provides a versatile method for characterizing EV subpopulations based on marker expression.

##### Preparation of Ab–Nb Conjugates

Anti-PSMA Ab–Nb labeled with Atto 488 or Alexa 647, and anti-CD33 Ab–Nb labeled with Atto 488 or Alexa 647, were generated by combining primary IgG anti-PSMA (absolute antibody, ab03136-23) or primary IgG anti-CD33 (Abcam, AB269456) with 2.Nb conjugated to either Atto 488 or Alexa 647. Reactions were set up in 0.5 mL Protein LoBind tubes (Eppendorf, cat. no. 0030108094) at a minimum volume of 5 µL. Both the primary IgG and 2.Nb were used at a final concentration of 50 nM, and the mixtures were incubated for 60 min at room temperature in the dark. Unless otherwise specified, samples were subsequently diluted as needed in 0.22 µm-filtered PBS (ThermoFisher Scientific, cat. no. 10010023).

##### Preparation of LnCAP-derived EVs

Lyophilized microvesicles derived from the LnCAP cell line (HBM-LnCAP-100, Hansabiomed) were reconstituted following the manufacturer's instructions, aliquoted into 20 µL portions, and stored at –80 °C. To minimize degradation, each aliquot was subjected to no more than one freeze-thaw cycle.

##### Complex Formation and Incubation with Extracellular Vesicles

Ab-Nb complexes composed of the same primary antibody but different nanobodies were prepared by mixing them in a 1:1 ratio. The complexes were then immediately incubated with various concentrations of LnCAP EVs at a final Ab–Nb concentration of 1 nM, at room temperature for 30 min in the dark. After incubation, the samples were fixed on ice for 15 min with 4% paraformaldehyde (PFA). Finally, the samples were diluted to 25 pM (Ab–Nb complex concentration) before subsequent analysis.

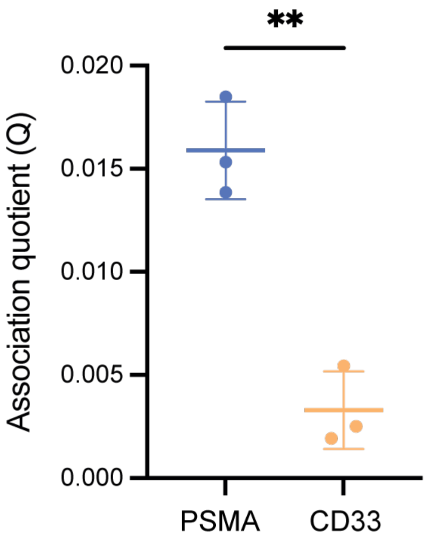

***Figure S4. Detection of LnCAP EVs with specific antibody (PSMA) or non specific antibody (CD33). Statistical significance was determined by paired two-tailed t-test.***

#### S6. Detection of EVs in biofluids

***Table S2 – mean intensities from donor serum and plasma.***

| | **Plasma** | | | **Serum** | | |
|---|---|---|---|---|---|---|
| **Sample** | Donor 1 | Donor 2 | Donor 3 | Donor 1 | Donor 2 | Donor 3 |
| **Mean intensity (photons bin$^{-1}$)** | 391.95 | 305.22 | 383.00 | 285.89 | 275.46 | 272.83 |

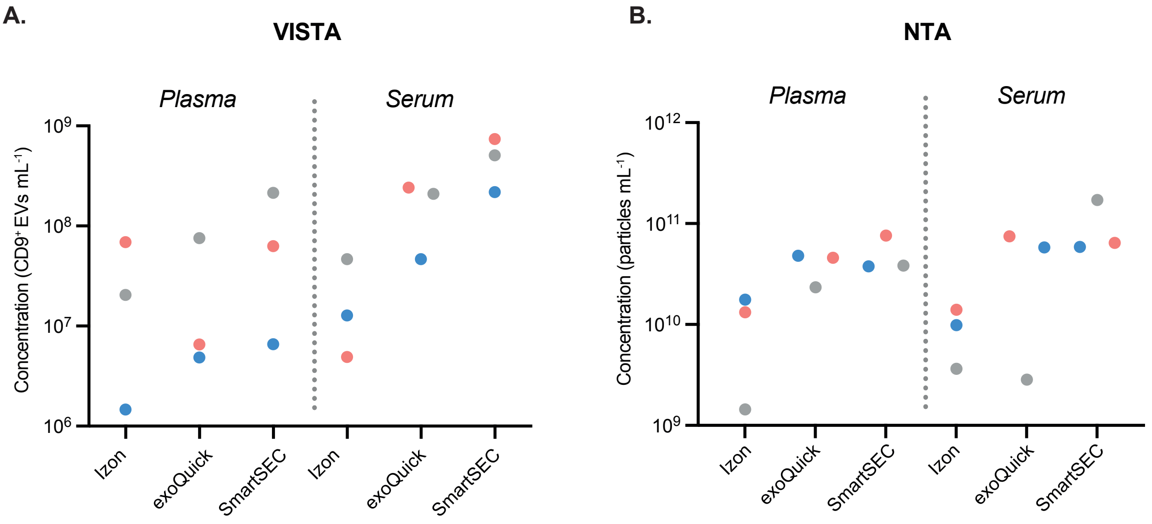

***Figure S5. Evaulation of different SEC isolation kits.*** *(A) VISTA measurements of Izon, exoQuick or SmartSEC (SEC) isolated EVs from plasma and serum, and (E) NTA measurements of same samples. The concentration of EVs has been normalized to the initial concentration from serum and plasma.*
